# Controlled enzyme cargo loading in engineered bacterial microcompartment shells

**DOI:** 10.1101/2024.10.21.619467

**Authors:** Nicholas M. Tefft, Yali Wang, Alexander Jussupow, Michael Feig, Michaela A. TerAvest

## Abstract

Bacterial microcompartments (BMCs) are nanometer-scale organelles with a protein-based shell that serve to co-localize and encapsulate metabolic enzymes. They may provide a range of benefits to improve pathway catalysis, including substrate channeling and selective permeability. Several groups are working toward using BMC shells as a platform for enhancing engineered metabolic pathways. The microcompartment shell of *Haliangium ochraceum* (HO) has emerged as a versatile and modular shell system that can be expressed and assembled outside its native host and with non-native cargo. Further, the HO shell has been modified to use the engineered protein conjugation system SpyCatcher-SpyTag for non-native cargo loading. Here, we used a model enzyme, triose phosphate isomerase (Tpi), to study non-native cargo loading into four HO shell variants and begin to understand maximal shell loading levels. We also measured activity of Tpi encapsulated in the HO shell variants and found that activity was determined by the amount of cargo loaded and was not strongly impacted by the predicted permeability of the shell variant to large molecules. All shell variants tested could be used to generate active, Tpi-loaded versions, but the simplest variants assembled most robustly. We propose that the simple variant is the most promising for continued development as a metabolic engineering platform.

## Introduction

Bacterial microcompartments (BMCs) have been discovered across many different species.^1^ BMCs consist of a protein shell and encapsulated enzymes. In the case of cyanobacteria, the carboxysome is a microcompartment that concentrates CO_2_ to enhance the carbon fixation efficiency of encapsulated ribulose-1,5-bisphosphate carboxylase/oxygenase (RuBisCo). In contrast with anabolic carboxysomes, metabolosomes are catabolic BMCs. The two most studied metabolosomes are the Eut BMCs which enable ethanolamine degradation and Pdu BMCs which are involved in 1,2-propanediol metabolism. Both are found in gut bacteria and are predicted to sequester toxic intermediates and improve enzymatic activity by co-localizing enzymes that produce and consume these intermediates.^2–5^

The BMCs identified from *Haliangium ochraceum* (HO) have a yet unknown function but have been recombinantly expressed in *Escherichia coli* and assembled both *in vivo* and *in vitro*.^6^ Five structural subunits are used in assembly of HO BMCs: BMC-H, BMC-T1, BMC-T2, BMC-T3, and BMC-P (hereafter, referred to as H, T1, T2, T3, and P). H is a hexamer that forms a hexagonal tile that forms the bulk of the structure. T1, T2 and T3 are trimers forming hexagonal tiles, with T2 and T3 trimers dimerizing to form stacked tiles while T1 remains unstacked. P is a pentamer that forms the vertices of the icosahedron. The various tiles that compose the HO shell can be used modularly to create a variety of shell types. Shells formed with only the T1-type of trimer are termed ‘minimal shells’ and BMCs containing all three trimers are described as ‘full shells’. Shells with P are called ‘capped’ shells and those without are ‘uncapped’ shells, also referred to as wiffle balls (**Figure 1**). By controlling which HO BMC subunits are present, we can create four distinct HO shell types: full-wiffle (HT1T2T3), full-capped (HT1T2T3P), minimal-wiffle (HT1), and minimal-capped (HT1P). These four shell types have the same size (∼40 nm) and icosahedral shape. ‘HP’ shells assembled with no trimer tiles have also been observed, although they are smaller (∼25 nm).^7^

**Figure 1.**
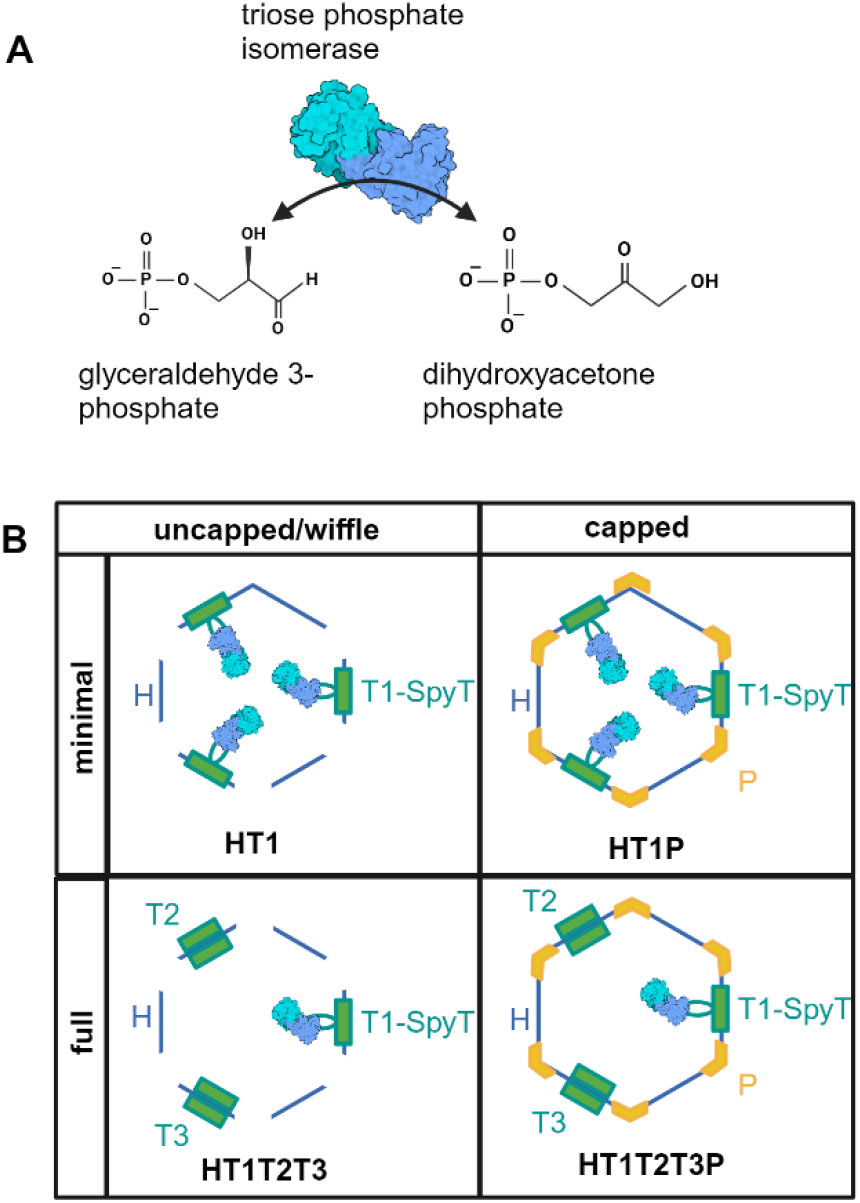
**A**. Illustration of Tpi structure (1TRE)^16^ and reaction. **B**. Cartoon representation of the four HO shell variants used in this study, included the tiles, cargo attachment points, and Tpi cargo. Labeled with terminology used throughout the text. Created in BioRender.com.

A key research goal is to understand how enzymatic cargo is localized to the BMC shell interior, both to understand encapsulation in natural systems and to enable loading of non-native cargo. Encapsulation peptides for Eut and Pdu BMCs have been identified and their roles in both native and non-native cargo loading have been determined.^8–12^ However, because the native encapsulation targets are not known for HO BMCs, encapsulation peptides are also unknown. For HO-BMCs an alternate strategy has been used, leveraging the strength of the SpyCatcher-SpyTag system to control the loading of targeted proteins for encapsulation.^13^ The SpyCatcher-SpyTag system consists of an engineered protein domain (SpyCatcher) and engineered tag (SpyTag) that spontaneously conjugate to form an isopeptide bond. In previous work, SpyTag was added to the interior surface of T1, allowing up to three molecules of protein cargo with SpyCatcher to be linked to each T1 tile.^13^

This localization system can create up to a three-fold difference in protein loading between full shells, which have ∼20 T1 subunits, and minimal shells, which have 60 T1 subunits. We can predict that with complete cargo loading, minimal shells will have increased enzyme crowding compared to full shells because T2 and T3 compete with T1 and do not localize protein cargo. Full cargo loading in a minimal shell could negatively impact shell assembly or enzyme function through steric hindrance. However, actual cargo loading is expected to be less than maximum, as observed previously.^14^ Similarly, we can predict that diffusion through uncapped shells (without P may be faster than diffusion through capped shells (with P), but this will be dependent on the target substrate and its interactions with the tile pores. The absence of a pentamer creates a much larger pore (∼6 nm) in the shell than is natively present in the T (∼2 nm) or H tiles (∼0.69 nm) that could enable faster diffusion rates across the shell boundary, especially for large or charged molecules.^14,15^

Understanding the effects of crowding and diffusion on encapsulated enzymes is essential to developing the HO BMC shells as a platform for encapsulating a range of enzymes and reactions. We have chosen triose phosphate isomerase (Tpi) as a model enzyme for understanding the effects of encapsulation in HO-BMCs on reaction rates. Tpi is a homodimer that is well-studied, stable, and of a compatible size for encapsulation within the HO-BMCs (27 kDa per monomer, 54 kDa per native enzyme). Further, activity can be measured using commercially available kits, enabling rapid evaluation of enzyme function within BMCs. Tpi is a glycolytic enzyme that catalyzes isomerization between dihydroxyacetone phosphate and glyceraldehyde 3-phosphate (**Figure 1**). To better understand the influence of enzyme cargo loading on assembly and the impacts of encapsulation on enzyme activity, we encapsulated *E. coli* Tpi in the HO shell system. We measured the levels of encapsulated cargo by both Western blot and untargeted proteomic analysis, and we assessed enzyme activity of free and encapsulated Tpi using a standard enzyme assay.

## Results

### Modeling Tpi loading into HO shells

We used modeling to estimate the theoretical maximum number of Tpi cargo molecules that could be encapsulated in an HO shell. We assumed that all Tpi in the shell are SpyCatcher-Tpi, i.e, there is no dimerization with native Tpi. The 50 amino acid N-terminal segment of the SpyCatcher001^17^ domain was omitted because it could not be reliably modeled. The modeling suggests that Tpi in the HO shell assembled into a quasi-symmetric pentameric framework that allowed loading of 30 Tpi dimers per shell (**Figure 2**). The SpyCatcher-SpyTag cargo loading system we used included the SpyTag on the T1 tile, and there are 20 total T tiles per shell. Therefore, the theoretical limit of cargo loading based on the conjugation system is 60 Tpi per minimal (T1 only) shell and 20 per full (T1, T2, and T3 shell). The modeling results suggest that at least for the minimal shell, conjugation of every SpyTag-T1 to SpyCatcher-Tpi would inhibit shell assembly.

**Figure 2.**
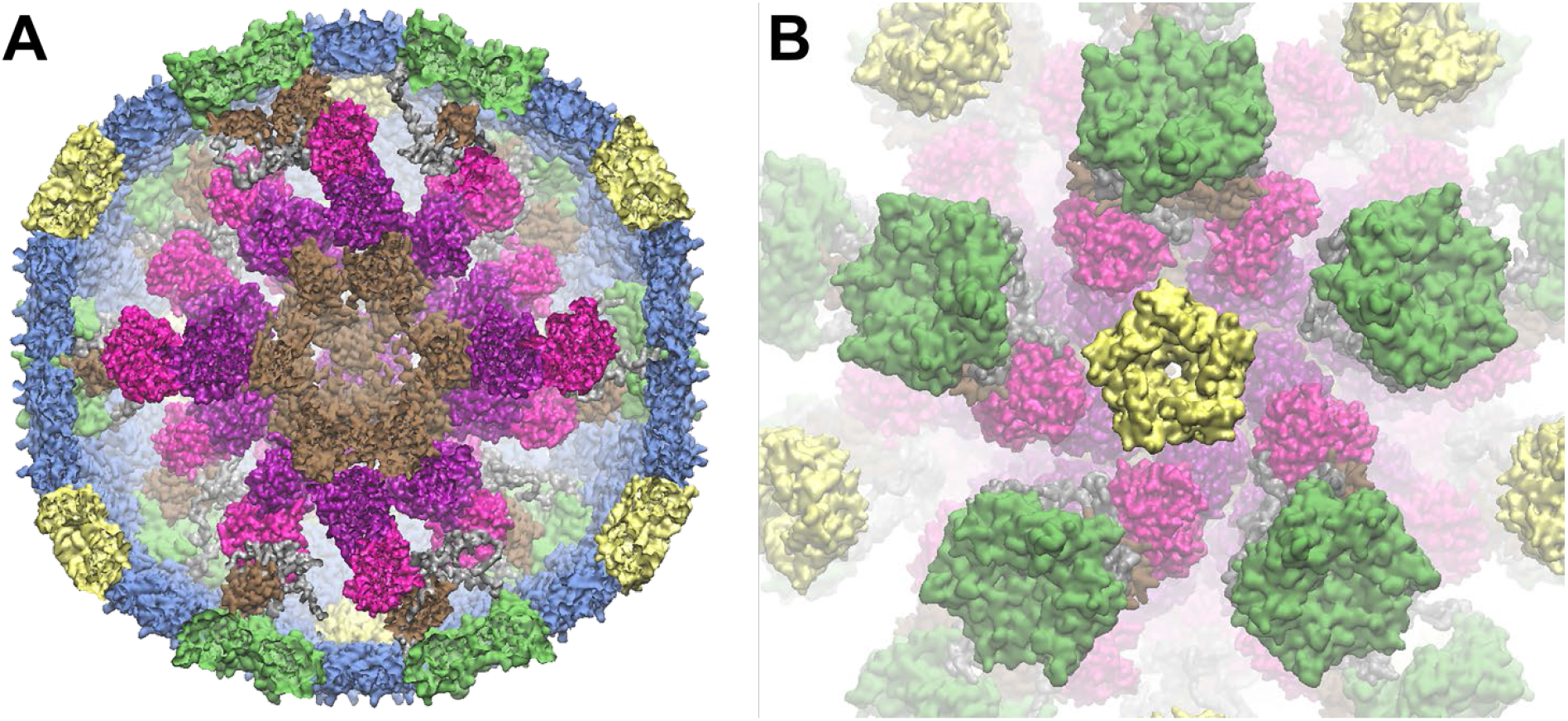
Model of Tpi-loaded HO shell. **A**. Cross-section of a Tpi-loaded minimal, capped HO shell. B. View into the HO shell from a P tile down, with H tiles omitted to enable viewing of Tpi cargo organization. TPI (magenta), SpyCatchers (brown), trimer (green), pentamer (yellow), hexamer (blue), SpyTag (grey).

### Design and purification of SpyCatcher-Tpi loaded HO shells

Experimentally,Tpi loading into HO shells was accomplished using the SpyCatcher-SpyTag system as previously described.^13,17,18^ To enable Tpi loading into HO shells, we created a Tpi-SpyCatcher fusion, with SpyCatcher (SpyC) added to the N-terminus of *E. coli* K-12 Tpi (NP_418354.1) with a glycine-serine linker. A Strep-tag II was also added to the N-terminus of the SpyC to enable purification of the modified Tpi independent of the HO shells. To ensure that SpyCatcher-Tpi retained Tpi activity, the fusion protein was purified using a StrepTrap HP column and was verified via sodium dodecyl sulfate-polyacrylamide gel electrophoresis (SDS-PAGE) (**Figure 3**). Activity of the purified SpyC-Tpi was confirmed using Tpi Activity Assay Kit (Abcam) (Figure S1). Genes encoding the modified Tpi and shell proteins were co-expressed in *E. coli* for production, assembly, and cargo-loading *in vivo* (**Table 1**). A SpyTag was previously inserted into an internal loop of the T1 trimer to enable conjugation of SpyCatcher fused cargo. Each shell variant was expressed from a different isopropyl β-D-1-thiogalactopyranoside (IPTG) inducible vector containing only the shell components needed for each variant. The ‘shell vectors’ were co-transformed into *E. coli* BL21(DE3) with an anhydrotetracycline (aTc)-inducible vector carrying SpyC-Tpi.

**Table 1.** Strains and plasmids used in this study.

| Strain |  | Description | Source |
| --- | --- | --- | --- |
| *E. coli* |  |  |  |
| BL21 (DE3) |  | Protein expression host | NEB |
| Plasmid | Vector | Description | Source |
| pARH360 | pBbA2K | SpyCatcher001-mTurquoise expression vector, aTc inducible vector. | 13 |
| pNT002 | pBbA2K | Modified pARH360, SpyCatcher001-Triose phosphate isomerase expression vector. StrepTagII purification tag. | This Study |
| pHK1 | pBbE6A | H_T1spyTsnoopT his_pStrep | Kerfeld lab |
| pHK2 | pBbE6A | H_T1spyTsnoopT_T2_T3 | Kerfeld lab |
| pHK3 | pBbE6A | H_T1spyTsnoopT his_T2_T3_pStrep | Kerfeld lab |
| pNT001 | pBbE6A | modified pHK1 with pStrep removed | This Study |

**Figure 3.**
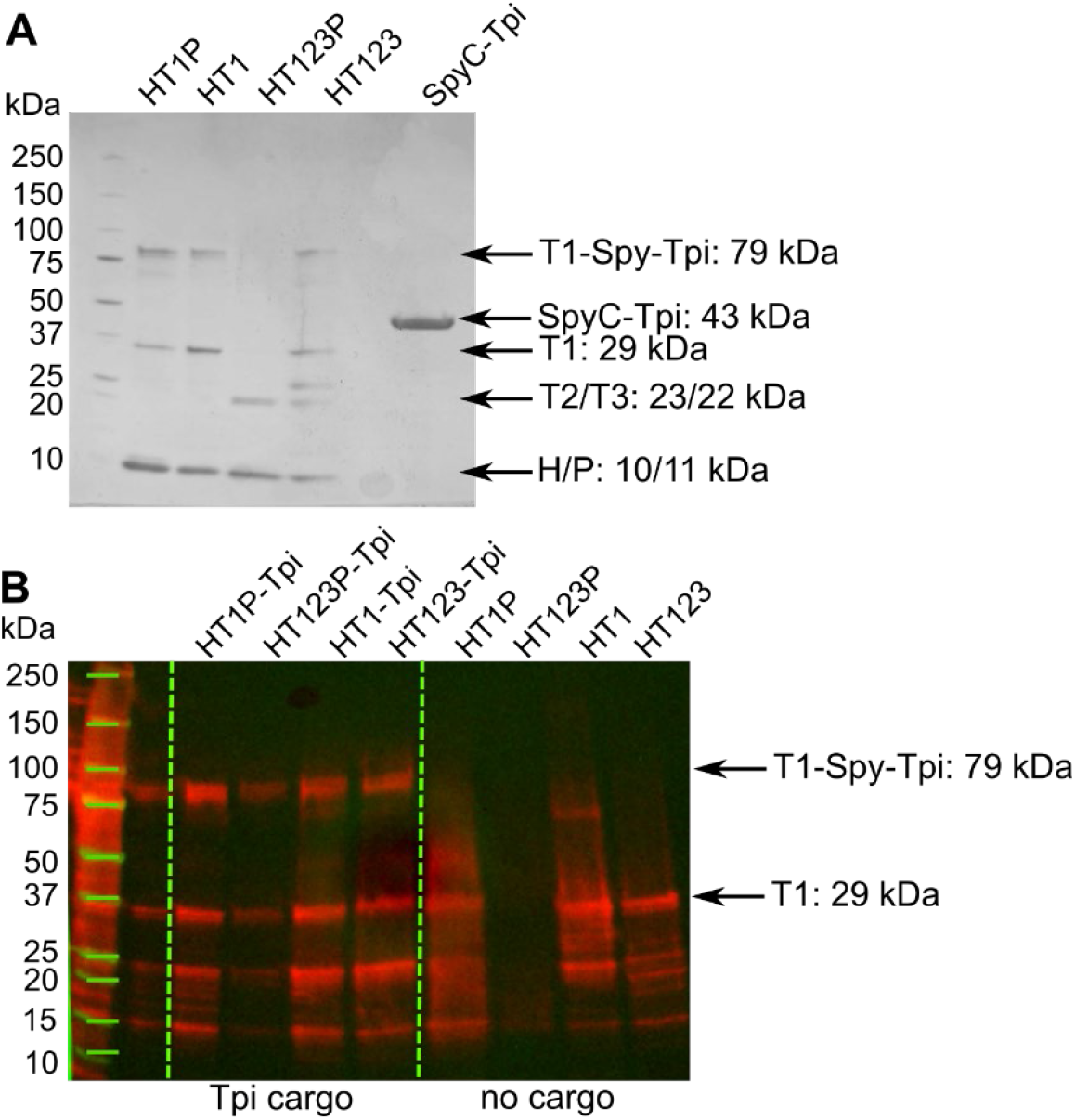
SDS-PAGE and Western blot of Tpi-loaded HO shells. **A**. Coomassie strained SDS-PAGE, 275 ng of protein loaded per well. **B**. Western blot of Tpi-loaded and empty HO shells using an anti-His antibody, 27.5 ng of protein loaded per well.

**Figure 4.**
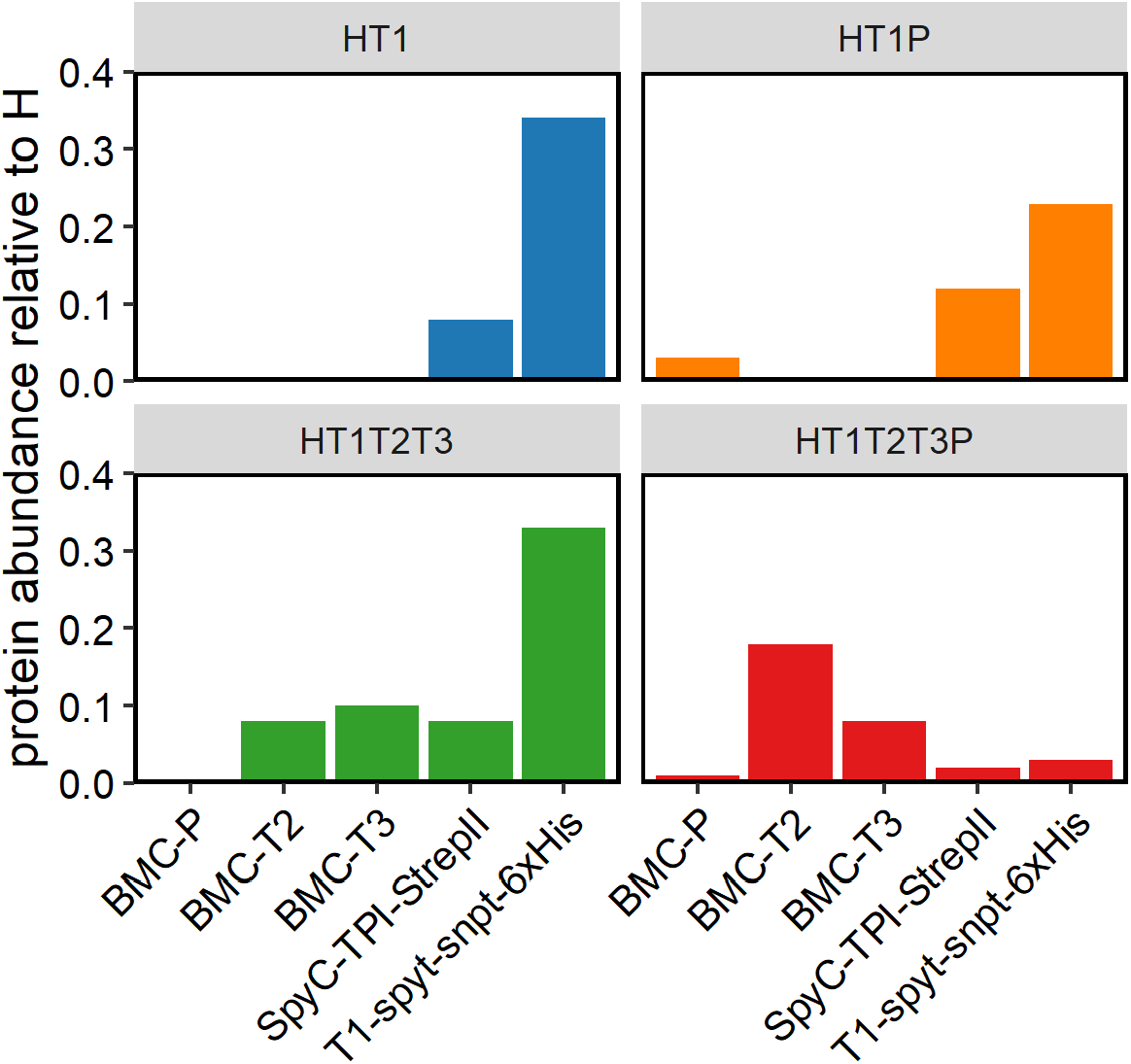
Untargeted proteomic analysis of cargo loaded HO shells. Purified cargo-loaded HO shell samples were submitted for untargeted proteomic analysis. Shell components and cargo signals were normalized to the signal for the hexamer. Error bars are not included because only one sample of each shell type was submitted for proteomic analysis.

Protein expression was induced for both vectors and shells were purified in a two-stage process using a His-Trap column against a His_6_-tag on the outer surface of the T1 followed by anion exchange chromatography. Each shell variant was also purified from *E. coli* without the ‘cargo vector’. Purified shells were visualized by SDS-PAGE (**Figure 3**). We observed assembly and purification of all four variants of the HO shell with Tpi; HT1P (minimal capped), HT1 (minimal wiffle), HT1T2T3P (full capped), HT1T2T3 (full wiffle). On SDS-PAGE, we observed the shell components expected for each sample, except the full capped shell, which appeared to lack T1 with or without conjugated cargo and T2. We did not observe any unconjugated SpyC-Tpi in any of the purified shell samples by SDS-PAGE. More sensitive detection by Western blot with anti-His_6_ antibodies to detect T1 and T1-Spy-Tpi confirmed the presence of both conjugated and unconjugated T1 in all shell types (**Figure 3**). This analysis also showed that the full capped shells did contain T1 with and without cargo, although not enough to be detected by standard Coomassie staining.

Further evaluation of shell loading was performed via untargeted proteomic analysis. All shell components and cargo were detected in each shell sample. We normalized the signal of other shell components to the signal of H, which is expected to be consistent across shell variants. Consistent with the Western Blot, less SpyC-Tpi was observed in HT1T2T3P shells than in other shell types. Further, HT1T2T3P, with and without TPI cargo, had less T1 and increased T2 and T3 compared with HT1T2T3. Other than HT1T2T3P, all other shell types show an abundance of unconjugated T1 in addition to T1-SpyT-SpyC-Tpi.

### Shell size and uniformity

We used transmission electron microscopy (TEM) to assess the shell size and uniformity for each shell type (**Figure 5**). The images show shells of the expected 40 nm size for all four shell types with varying impurities in each. HT1P shells show some smaller species, possibly HP shells based on relative size compared to the 40 nm species. HP shells may arise due to the inability for all BMC-T1-SpyCatcher-TPI to be incorporated into shells, leaving excess H unincorporated.^6^ The HT1T2T3P shell sample also contains a spindle shaped protein of unknown composition. It is difficult to predict what effect these proteins may have on Tpi activity.

**Figure 5.**
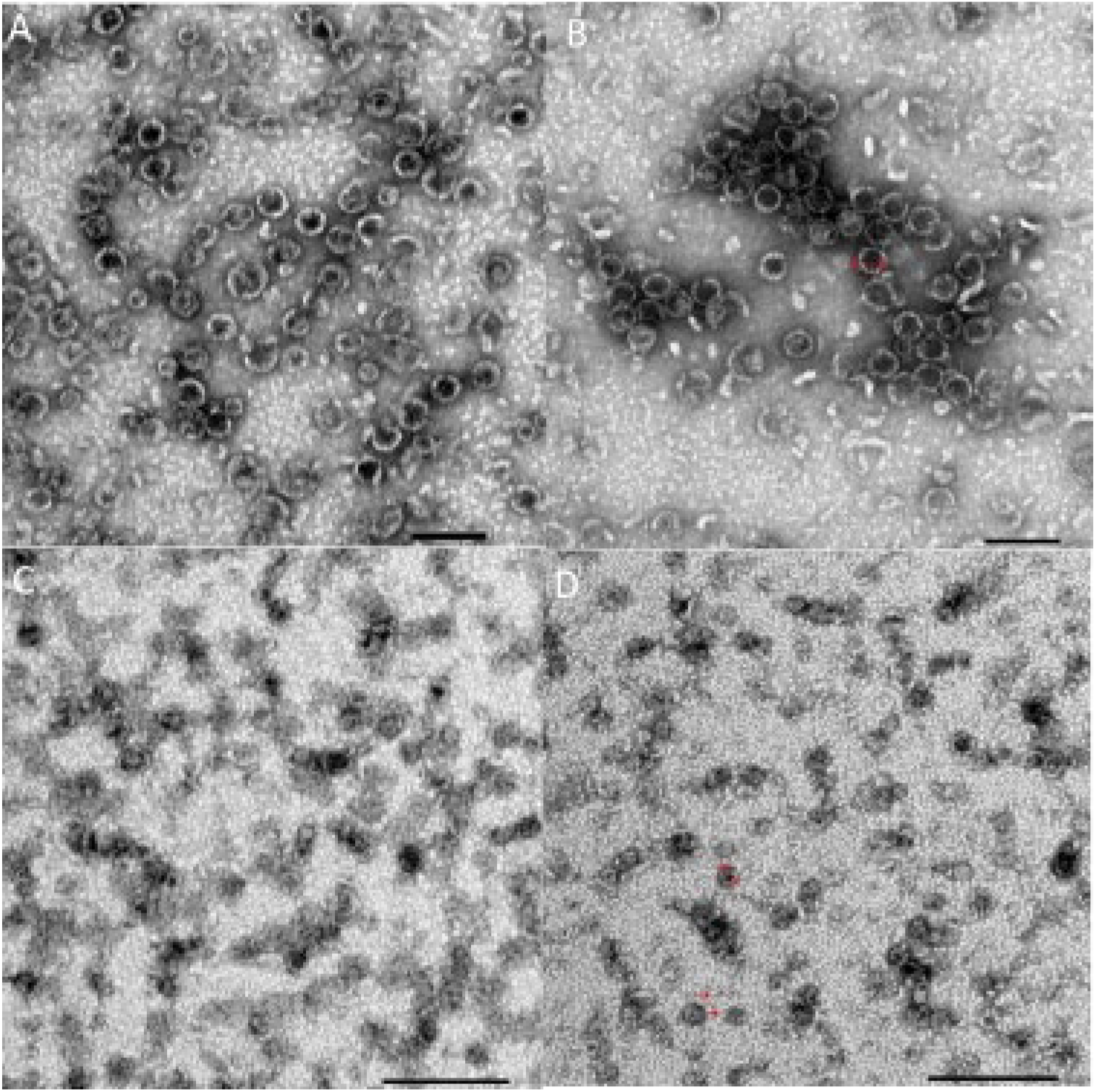
Transmission electron microscopy images of TPI BMCs. **A**. HT1P **B**. HT1T2T3P **C**. HT1 **D**. HT1T2T3

Both uncapped shell samples show similar images under TEM with shells joined by what appear to be imperfectly assembled complexes and non-uniform large complexes. Dynamic light scattering analysis (DLS) also indicated shell assembly and uniformity for all samples, with the average particle diameter ranging from 37 nm to 49 nm (Table S1).

### Tpi activity

After confirming shell assembly and cargo loading, we measured Tpi activity of all four shell types with and without cargo. We observed that all Tpi-loaded shell samples had much higher Tpi activity than their empty counterparts (**Figure 6**). All empty shell samples showed similar, low Tpi activity, which may be caused by low levels of sample contamination with native Tpi from the *E. coli* host. HT1P shells had higher Tpi activity per mg protein than any other shell type. We observed that the uncapped HT1 shells had slightly lower activity than the capped shells, suggesting that capping does not significantly alter the diffusion rate of DHAP or G3P across the shell boundary. Both capped and uncapped full shell variants had lower Tpi activity than the T1 only variants, consistent with the hypothesis that these would encapsulate fewer Tpi. HT1T2T3P shells had much lower Tpi activity than any other variant, as expected given that Tpi encapsulation appeared lower by Western blot and proteomic analysis.

**Figure 6.**
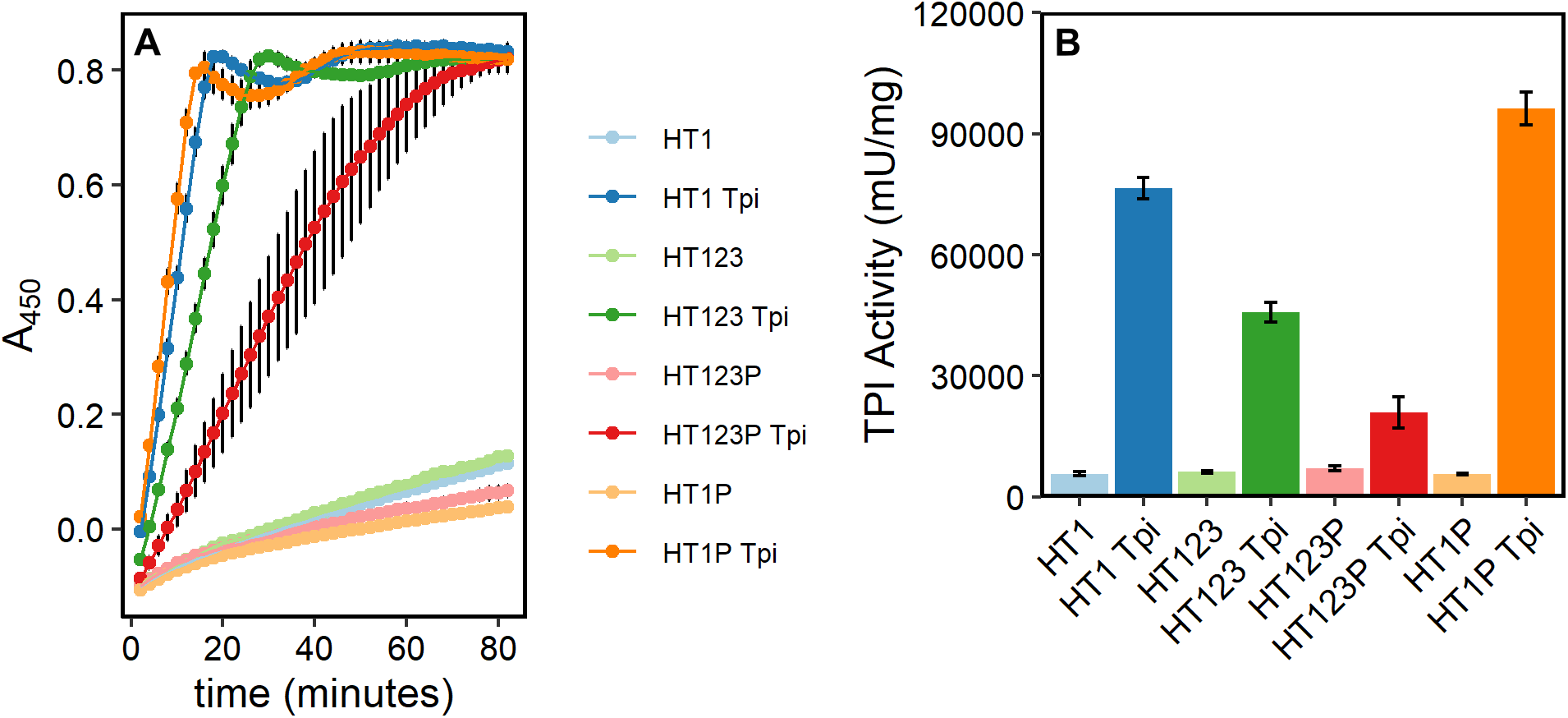
Tpi Activity of cargo-loaded HO shells. **A**. Increase in absorbance at 450 nm (A_450_) over time in a Tpi activity assay with 238 ng protein loaded per well for each sample. Each point represents the average of three replicates with standard deviation shown in error bar. HT1T2T3P is an average of 6 replicates. **B**. Calculated Tpi activity per mg protein from results in panel A.

To determine whether capped and uncapped shells had similar Tpi activity due to incomplete capping, we repeated the activity assay in the presence of excess P protein that was purified separately. We observed no difference in Tpi activity between samples with excess P, regardless of whether they were initially capped (**Figure 7**). We also added excess P to shells without cargo and observed no change in activity, indicating that the purified P protein was not contaminated with Tpi.

**Figure 7.**
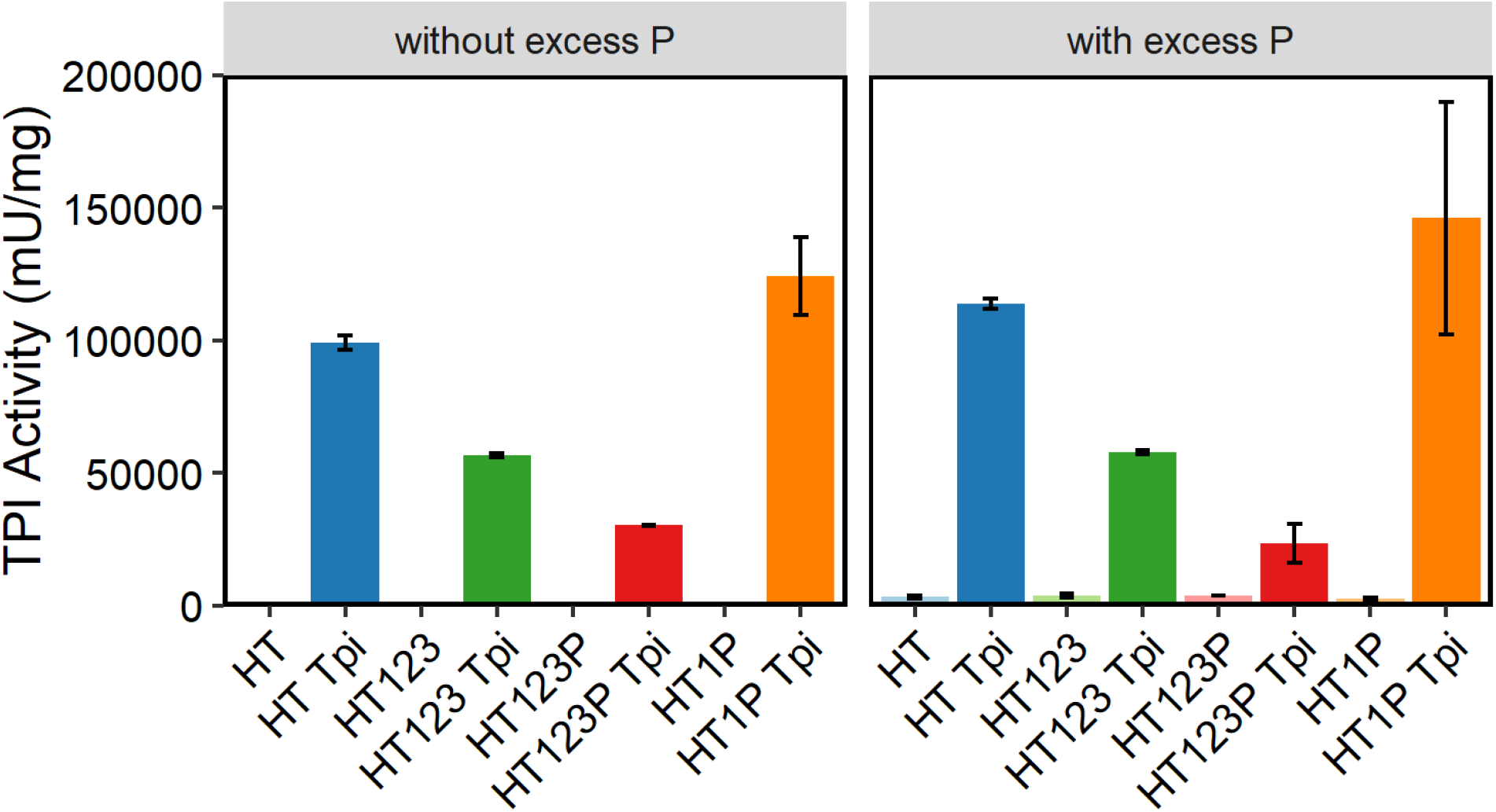
Tpi activity with and without excess P in a Tpi assay with 238 ng protein loaded per well for each sample. On the right, excess P was also added. Shell samples without cargo were not tested without excess P in this experiment. Each point represents the average of three replicates with standard deviation shown in error bar.

## Discussion

Engineered bacterial microcompartment systems hold the promise of enhancing catalysis in metabolic pathways by concentrating reaction substrates and protecting cellular processes from harmful intermediates. To realize this potential, it is necessary to control assembly and properties of the engineered systems, including enzyme encapsulation and permeability across the shell boundary. The BMC shell system from *Haliangium ochraceum* was identified through homology analysis and has proven to be a robust and modular BMC shell system for modification and expression in other bacterial hosts.^6,14,19^ The modular nature of the HO shell enables some aspects of cargo loading and permeability to be tuned simply by expressing a different subset of shell tiles. For example, we hypothesize that permeability for some molecules can be increased by omitting the P tiles and that cargo loading can be increased by omitting T2 and T3 tiles, which do not contain a SpyTag for cargo conjugation in the current engineered platform. Other researchers recently observed that capping influences the permeability of a large, positively charged molecule.^20^

Here, we investigated cargo loading by measuring loading efficiency, shell assembly, and enzyme activity of a model enzyme (Tpi) in four different HO shell variants. We successfully isolated Tpi-loaded versions of all four variants, confirming cargo loading by SDS-PAGE, Western blot, proteomic analysis, and enzyme activity. We observed significant differences between the shell types in terms of their ability to assemble with Tpi cargo. The ‘minimal shell’ variants with only the T1 trimer assembled more robustly and with higher cargo loading efficiency than ‘full shell’ variants with all three trimer types. However, we observed that not all T1 tiles in the minimal shells were conjugated to cargo, suggesting that if every T tile was conjugated, assembly could be impacted. Indeed, when cargo expression was increased with greater concentrations of aTc, we were not able to purify assembled shells from *E. coli* (data not shown). The SDS-PAGE and Western blot analysis both suggest similar levels of conjugated and unconjugated T1 tiles, which aligns well with modeling results suggesting that maximal packing would allow 30 Tpi per shell, or half of the 60 T1-SpyTag being conjugated to Tpi cargo.

As additional evidence that too much cargo could impede assembly, we observed that full shells incorporated more T2 and T3 tiles, suggesting that Tpi-conjugated T1 tiles were less preferred. Interestingly, overrepresentation of T2 and T3 tiles was more prominent in the capped shell variant than in the uncapped variant, indicating that the presence of the P tile alters interactions or assembly in a way that affects cargo loading efficiency. Overall, we find that controlling production of the cargo protein via inducible expression was a more successful strategy to optimize cargo loading and assembly than using different combinations of shell proteins. In this study, it was not possible to specifically determine the level of T1 conjugation that impacted shell assembly because high aTc concentrations yielded a lack of purified protein. However, in the future, a detailed investigation of the percentage of allowable conjugation for the T tiles could be conducted by *in vitro* assembly, where it would be possible to directly control the levels of conjugated and unconjugated T tiles.

Interestingly, we did not observe any systematic difference in activity between capped and uncapped shell variants, indicating that permeability of the shell to G3P and DHAP was not affected by the presence of P tiles. We repeated this experiment in the presence of excess P tiles to ensure that the result was not caused by missing P tiles in the capped shell samples. Previous work with the HO shell system has indicated that *post hoc* addition of P tiles can cap the shells and block permeation of a large, positively charged molecule.^20^ Addition of excess P tiles had no impact on Tpi activity, suggesting that capping did not create a diffusion barrier to the molecules of interest. We speculate that G3P and DHAP are small enough to pass through pores in the T tiles. This is unsurprising, given that other microcompartment systems, such as Pdu, process metabolites of similar size; 1,2-propanediol is also a 3-carbon molecule.^21^

Overall, our results indicate that HO shells likely cannot assemble with enzyme cargo conjugated to every T1 subunit, but that cargo loaded shells can readily be purified by expressing lower levels of the cargo enzyme. This approach was more successful than reducing cargo loading by expressing T2 and T3 tiles to reduce the number of SpyTag sites for conjugation. Modeling aligned with experimental outcomes, suggesting that 50% cargo loading leads to tight packing of the shell interior and the greater cargo loading efficiency may be unlikely. Future studies with *in vitro* assembly will shed light on the exact ratio of conjugated to unconjugated T1 that allows assembly.

## Materials and Methods

### Bacterial Strains, Plasmids, and Growth Conditions

Strains and plasmids used in this study are listed in Table 1. pNT001 was generated via PCR amplification of pHK1 omitting Hoch_5814 (BMC-P) and ligated using T4 ligase (NEB) after purification using a Qiaquick Gel Extraction Kit (Qiagen). *E. coli* BL21(DE3) chemically competent cells (Thermo Scientific) were transformed with 15-20 µg/ml of target plasmid(s) using the manufacturer’s protocol. After overnight growth, single colonies from the transformation were grown overnight in lysogeny broth (LB) (Miller, Fisher) shaking at 250 rpm at 37°C. Antibiotics were used at the following concentrations: 100 µg/ml ampicillin and/or 50 µg/ml kanamycin. For protein purification, 1.5 l of LB was inoculated to an OD of 0.01 and induced to a final concentration of 100 µM IPTG (GoldBio) and/or 50 ng/ml tetracycline (Sigma) before growth overnight at 30°C for protein expression.

### Protein purification

3 l of cells were harvested after 16-18 h of growth by centrifugation at 8000 x g for 10 min at 4°C (Sorvall LYNX 6000, Thermo Fisher) and resuspended in 100 ml of 50 mM Tris pH 8.0, 50 mM NaCl, and 20 mM imidazole by vortexing. 400 µl of DNAse I (Millipore Sigma) and 1 tablet of SigmaFast protease inhibitor (Millipore Sigma) was added. Resuspended cells were lysed by passage through a pre-cooled French press at 1,100 PSI twice. Lysate was clarified by centrifugation at 45,000 x g for 30 min at 4°C (centrifuge name and brand). Supernatant was removed and filtered through a 0.22 µm syringe filter. Proteins were purified using a ÄKTA pure fast protein liquid chromatography (Cytiva) and a 5 ml HisTrap (Cytiva) and eluted with 50 mM Tris pH 8.0, 50 mM NaCl and 300 mM Imidazole. BMC proteins were filtered using a 0.22 µm syringe filter before being further purified using a 5/50 GL MonoQ column (Cytiva) with a NaCl gradient to separate assembled shells from unassembled cargo and shell components.

Samples were eluted with 50 mM Tris pH 8.0, 1M NaCl, with intact shells appearing in 40-42% NaCl fractions.

For SpyCatcher-TPI protein alone a StrepTrap HP column (Cytiva) was used and eluted with 50 mM Tris pH 8.0, mM NaCl and 2.5 mM desthiobiotin.

Empty pHK1, pHK2, NT002, and cargo loaded pHK2+SpyCatcher TPI BMCs were concentrated using 100 kDa Amicon centrifugal filters (Millipore Sigma).

### Triose phosphate isomerase activity assay

Tpi activity was measured using Triose Phosphate Isomerase (TPI) Activity Assay Kit (Colorimetric) (ab197001) as described in the product manual. Protein samples were diluted prior to assay to normalize to total protein.

### SDS-PAGE

Protein concentration was measured via bicinchoninic acid (BCA) assay (Thermo Fisher) and samples were normalized to ensure consistent protein loading. 10 μl of each normalized sample was heated at 95°C for 10 min in 10 μl of a mixture of 1 ml 5x Laemmli buffer, 20 μl concentrated bromophenol blue (JT Baker, D29303) in 5x Laemmli buffer, and 10 μl of 1 M DTT. A mini-PROTEAN tetra cell electrophoresis chamber (Biorad, 1658005EDU) was loaded with 1x TGS buffer. 20 µl was loaded onto a mini-protean TGX stain free gel (Bio-Rad, 4568095) alongside 5 μL of Precision Plus Unstained ladder (Biorad, 1610363). Samples were run at 300 V for 20 min until the dye front moved off the gel. Gels were stained using Coomassie blue for 30 min, then destained in 10% methanol, 10% acetic acid overnight before imaging.

### Western Blot

Western blotting was performed as above for SDS-PAGE but the gel was removed to 1x Transfer buffer (Bio-Rad, 10026938) after electrophoresis. Proteins were transferred to a nitrocellulose membrane (Biorad, 1704270) using a Bio-Rad TurboBlot transfer system (Bio-Rad, 1704150). The membrane was rinsed with 30 ml Tris buffered saline with Tween 20 (TBST); this buffer was discarded, and the membrane was blocked using 50 ml of 3% bovine serum albumin (BSA) in TBST for 1 h on an orbital shaker. Blocking solution was discarded and replaced with 50 ml of 3% BSA TBST buffer, and 10 μL of anti-His_6_ antibody (GeneScript, 6G2A9) was added. The membrane was incubated for 16 h at 4°C on an orbital shaker.

The membrane was rinsed for 5 min with TBST three times. After rinsing, 50 ml of 3% BSA TBST buffer with 0.0625 μL of anti-mouse antibody (Sigma-Aldrich, A9044) was added. The membrane was incubated for 2 h at room temperature (RT). Buffer was discarded and the membrane rinsed with TBST buffer for 5 min, three times. ECL Clarity chemiluminescence solution (Bio-Rad, 1705061) was prepared by mixing 10 ml of peroxide solution with 10 ml of enhancer solution and adding the entire volume to the membrane. The membrane was incubated for 5 min; the ECL solution was discarded, and the membrane was imaged using a Bio-Rad Molecular Imager Gel Doc XR.

### Dynamic light scattering analysis

Dynamic light scattering was performed on a Wyatt DynaPro (Nanostar). 10 μL of the HO shell samples were centrifuged for 5 min at 13,000 x g before being loaded into 1×1×10 mm cuvette. Samples were scanned 20 times with 5 second acquisitions. This was repeated three times on each sample to measure shell diameter.

### Transmission electron microscopy

For negative-staining TEM, 10 µl of purified protein was placed on a Formvar/Carbon 200 mesh Cu grid and incubated for 1 min. The grid was washed twice with 15 µl of ddH_2_O and blotted with filter paper. It was stained with 1% (w/v) uranyl acetate for 40 s before being blotted nearly dry. The prepared grid was imaged using a JEOL 1400 Flash transmission electron microscope at an operating voltage of 100 V.

### Proteomic analysis

Samples were mixed with 4% (w/v) sodium deoxycholate (SDC) in 100 mM Tris, pH 8.5, reduced and alkylated by adding Tris(2-carboxyethyl)phosphine (TCEP) and iodoacetamide at 10 mM and 40 mM, respectively, and incubated for 5 min at 45°C with shaking at 2000 rpm in an Eppendorf ThermoMixer R. Trypsin/LysC enzyme mixture, in 50 mM ammonium bicarbonate, was added at a 1:100 ratio (wt/wt) and the mixture was incubated at 37°C overnight with shaking at 1500 rpm in the ThermoMixer. Final volume of each digest was ∼300 µl. After digestion, SDC was removed by adding an equal volume of ethyl acetate and trifluoracetic acid (TFA) to 1% (v/v). Samples were centrifuged at 16,800 x g for 3 min to pellet SDC and separate the aqueous and organic phases. The aqueous phase was removed to a new tube and dried briefly by vacuum centrifugation to remove residual ethyl acetate. Peptides were subjected to C18 solid phase clean up using StageTips^1^ to remove salts and eluates and dried by vacuum centrifugation.

Dried peptides were re-suspended in 20 µL of 2% acetonitrile/0.1% TFA. The sample was diluted 1:10 on plate and an injection of 2 µL was automatically made using a Thermo EASYnLC 1200 onto a Thermo Acclaim PepMap RSLC 0.1mm x 20mm C18 trapping column and washed for ∼5 min with buffer A. Bound peptides were eluted over 35 min onto a Thermo Acclaim PepMap RSLC 0.075 mm x 250 mm resolving column with a gradient of 5% B to 19% B from 0 min to 19 min and 19% B to 40% B from 19 min to 24 min (Buffer A = 99.9% water/0.1% formic acid, Buffer B = 80% acetonitrile/0.1% formic acid/19.9% water) at a constant flow rate of 300 nl/min. After the gradient the column was washed with 90% B for the duration of the run. Column temperature was maintained at a constant temperature of 50°C using and integrated column oven (PRSO-V2, Sonation GmbH, Biberach, Germany).

Eluted peptides were sprayed into a ThermoScientific Q-Exactive HF-X mass spectrometer using a FlexSpray spray ion source. Survey scans were taken in the Orbi trap (60,000 resolution, determined at m/z 200) and the top 10 ions in each survey scan were subjected to automatic higher energy collision induced dissociation (HCD) with fragment spectra acquired at a resolution of 15,000. The resulting MS/MS spectra were converted to peak lists using Mascot Distiller (www.matrixscience.com), v2.8.5 and searched against a protein sequence database containing all entries for *E. coli* (downloaded from www.uniprot.org on 2022-11-30) appended with customer provided sequences and common laboratory contaminants (downloaded from www.thegpm.org, cRAP project) using the Mascot^2^ search algorithm, v2.8.3. The Mascot output was analyzed using Scaffold, v5.3.3 (www.proteomesoftware.com), to probabilistically validate protein identifications. Assignments validated using the Scaffold 1% FDR confidence filter are considered true. Mascot parameters for all databases were as follows: allow up to 2 missed tryptic sites; fixed modification of carbamidomethyl cysteine; variable modification of oxidation of methionine; peptide tolerance of +/-10 ppm; MS/MS tolerance of 0.02 Da; false discovery rate (FDR) was calculated using randomized database search.^22,23^

### Computational modeling

Cargo-loaded shells were constructed starting from the cryo-electron microscopy structure of a complete HO BMC shell (PDB code: 6MZX).^24^ T2 trimers in this structure were replaced by T1 trimers via superposition by using the T1 structure from PDB code 5DIH.^25^ A SpyCatcher-SpyTag domain was then modeled according to the structure deposited under PDB code 4MLI. The SpyCatcher-SpyTag complex was initially positioned near a T1 subunit with enough space for a flexible linker to connect to residue 84. The linker was modeled in random conformation with amino acids mutated to match the sequence in the experimental construct. The experimental construct also includes a SnoopTag which was modeled according to the structure in PDB code 2WW8,^26^ but without the SnoopCatcher. The SnoopTag sequence was manually placed together with flexible linkers to avoid the initially placed SpyCatcher-SpyTag complex while connecting back to the insertion site in the T1 subunit. The SpyCatcher domain was then extended with a Tpi domain using the structure from PDB code 1TRE.^16^ Tpi was modeled as a dimer with the dimer interface according to the crystal structure and with another SpyCatcher domain attached to the second dimer moiety facing away from the T1 structure. The model for one T1 subunit with attached SpyTag:SpyCatcher and a Tpi dimer was replicated for all T1 subunits by superimposing the T1 structure from one subunit onto one of the other subunits. In the initial placement, the SpyCatcher and Tpi domains were relatively far from T1 to avoid structural overlap. However, the initial model was too extended to allow placement into the HO shell without clashes, requiring relaxation of the initial model into a more compact arrangement. This was accomplished via minimization under restraints that kept the T1 trimer fixed in space and allowed only rigid body motions of the folded domains (SpyCatcher and Tpi) while leaving the flexible linkers unrestrained. A bias was then applied during step-wise minimization runs to reduce the distances between the T1, SpyCatcher, and Tpi domains to achieve close packing. During minimization, all domains were modeled in atomistic detail and a distance-dependent dielectric was applied to mimic solvation effects. The modeling and minimization runs were carried out with the MMTSB Tool Set^27^ in combination with CHARMM.^28^ VMD^29^ was used for visualization and for the initial manual placements.

## Acknowledgements

Research was primarily supported as part of the Center for Catalysis in Biomimetic Confinement, an Energy Frontier Research Center funded by the U.S. Department of Energy (DOE), Office of Science, Basic Energy Sciences (BES), under award DE-SC0023395. Additional support from National Institutes of Health (R35 GM126948, to MF) is acknowledged. The proteomic analysis was performed with Douglas Whitten and the MSU Proteomics Core Facility. We thank Matthew Dwyer for training on equipment and troubleshooting protein purification protocols

## Notes

### Competing Interest Statement

The authors have declared no competing interest.

