## Supplementary information for "Controlled enzyme cargo loading in engineered bacterial microcompartment shells"

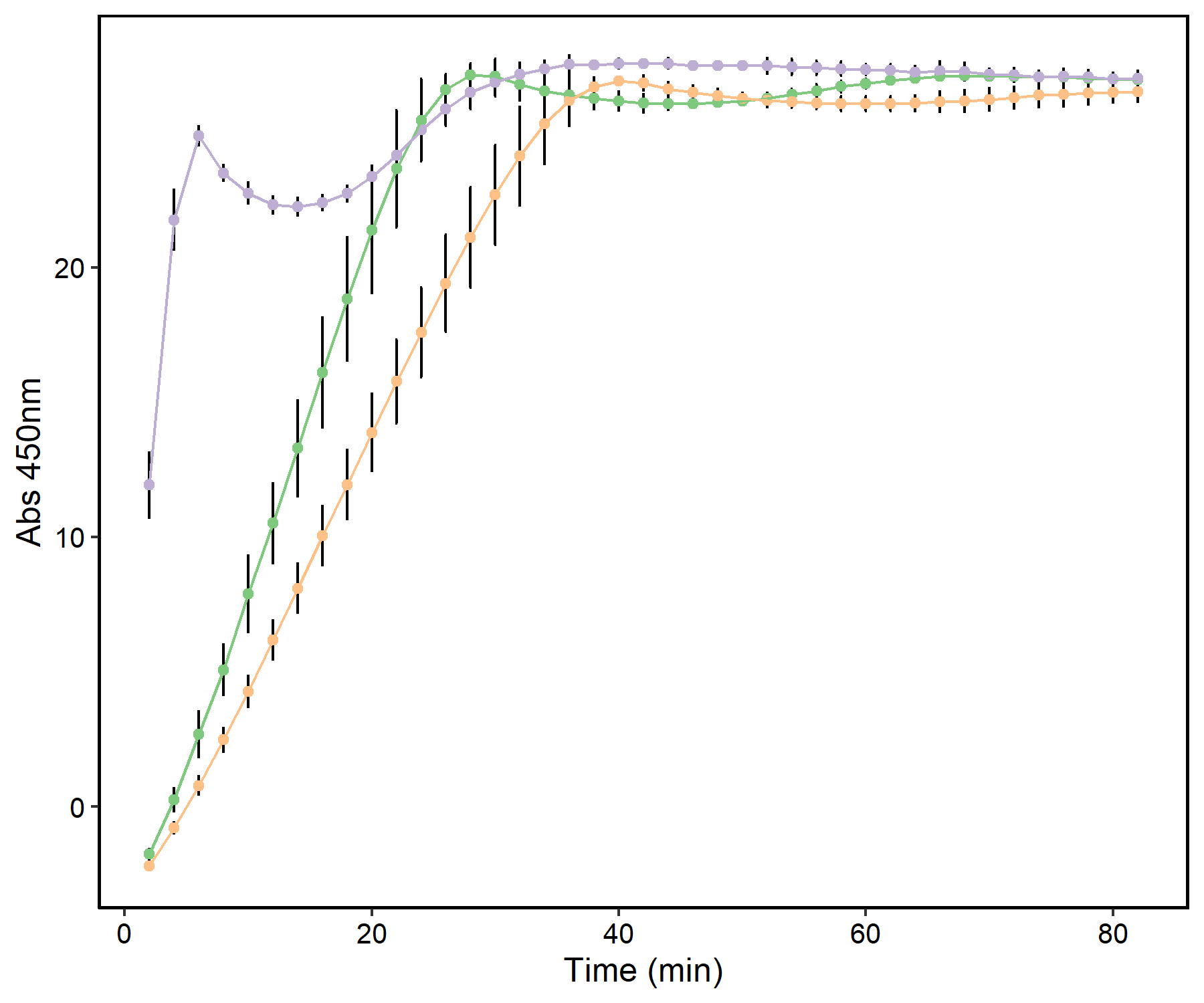

**Figure S1**. **TPI activity of purified SpyCatcher-TPI.** Purple: 238 ng SpyCatcher TPI. Green: TPI positive control . Orange: 23.8 ng SpyCatcher TPI.

**TableS1.** Dynamic Light Scattering analysis of shell samples.

| Shell | Average diameter | stdev |
| --- | --- | --- |
| HT1P | 37.6 | 1.6 |
| HT1T2T3P | 39.7 | 1.8 |
| HT1 | 49.1 | 1.5 |
| HT1T2T3 | 43.2 | 0.8 |
